# Comparative Analysis of Pathology Foundation Models for Automated Detection of Tertiary Lymphoid Structures in H&E-Stained Digital Pathology Images

**DOI:** 10.1101/2025.11.23.688074

**Authors:** Meijian Guan, Yu Sun, Merzu Belete, Anantharaman Muthuswamy, Maximilian Farma, Jenny Kaufmann, Mirna Lechpammer, Sriram Sridhar, Brandon W. Higgs, Han Si

**Affiliations:** Translational Data Science, Genmab, Princeton, NJ, USA; Pathology, Genmab, Princeton, NJ, USA; Cornell University, Ithaca, NY, USA; Harvard University, Cambridge, MA, USA

**Author notes:** Correspondence to: Meijian Guan and Han Si Building 2, 777 Scudders Mill Rd, Princeton, NJ 08540.

## Abstract

Tertiary lymphoid structures (TLS) have been observed in solid tumors and have been associated with better outcomes in patients treated with immunotherapy, but their dynamic nature makes identifying TLS in clinical samples challenging. Recently, pathology foundation models have emerged as powerful tools in computational pathology. In this study, we aimed to develop a computational tool capable of identifying TLS across different cancer types. To this end, we utilized multiple pathology foundation models, along with the ImageNet-pretrained ResNet50 as a baseline, to identify TLS in pancreatic ductal adenocarcinoma (PDAC) and head and neck squamous cell carcinoma (HNSCC) using hematoxylin and eosin (H&E) stained images from The Cancer Genome Atlas (TCGA) and a licensed Real-World Evidence (RWE) cohort. Both pathologist-annotated and transcriptomic signature-based TLS outcomes were employed for performance assessment. Pathologist-identified TLS-positive tumors showed higher expression of TLS signatures in both diseases.

Among the models tested, PLIP and CTransPath exhibited strong performance in identifying TLS within PDAC samples (AUC = 0.94 and 0.89, respectively). However, all models struggled in the analysis of HNSCC, likely due to the increased heterogeneity of the tumor microenvironment (TME).

Despite their overall utility in detecting TLS in PDAC, all foundation models demonstrated poor performance in predicting transcriptomic signature-based outcomes in both PDAC and HNSCC. This suggests that TLS signatures may reflect broader or more transient aspects of TLS biology, whereas pathology-based assessment anchors on visible morphological features, more closely aligned with the focus of foundation model training.

These findings highlight the potential of advanced pathology foundation models for TLS detection and broader tumor immune profiling tasks. These models can also be utilized on routinely collected patient biosamples with nominal costs. However, further refinement is needed to enhance their utility in tumors with more complex TME. Additionally, the identification of transcriptomics-based biomarkers from H&E images remains a significant challenge, despite advancements in digital pathology.

## Introduction

Tertiary lymphoid structures (TLS)^1^ are ectopic lymphoid aggregate structures of immune cells that form within non-lymphoid tissues, including in the tumor microenvironment^2^. Found in various cancers, including head and neck squamous cell carcinoma (HNSCC)^3^ and pancreatic ductal adenocarcinoma (PDAC)^4, 5^, TLS are composed of B-cell follicles, T-cell zones and dendritic cells, and function as sites for local immune activation.

TLS are crucial in cancer immunotherapy due to their role in modulating anti-tumor immune responses^6-8^. They act as hubs for antigen presentation, T-cell activation, and B-cell maturation, fostering a robust immune response against tumor cells. For example, TLS presence is associated with better response rates to immune checkpoint inhibitors, as they provide a localized immune environment that enhances T-cell activation and cytotoxic activity. Consequently, TLS are emerging as predictive biomarkers for immunotherapy efficacy and overall survival in cancer patients.

Recent studies have highlighted the clinical significance of TLS in various cancers. A meta-analysis encompassing 17 studies with 4,291 lung cancer patients revealed that a high TLS presence correlates with improved overall survival (HR = 0.66, 95% CI: 0.50–0.88) and disease-free survival (HR = 0.46, 95% CI: 0.33–0.64)^9^. Similarly, research focusing on digestive system cancers demonstrated that the absence of TLS is associated with poorer overall survival (HR = 1.74, 95% CI: 1.50–2.03) and recurrence-free survival (HR = 1.96, 95% CI: 1.58–2.44)^10^.

Furthermore, the presence of mature TLS has been linked to enhanced responses to immune checkpoint inhibitors, suggesting their potential as predictive biomarkers for immunotherapy efficacy^11^. In HNSCC, intratumoral TLS have been associated with improved patient survival and better responses to immunotherapy^12^. Similarly, research indicates that patients with PDAC who exhibit TLS tend to have a more favorable prognosis compared to those who do not^5, 13, 14^. Collectively, these findings highlight the prognostic and therapeutic relevance of TLS across multiple cancer types.

TLS detection has traditionally relied on RNA sequencing (RNAseq) to identify specific gene signatures^3, 15, 16^, manual histology-based assessment to visually identify structures within tumor tissues^17, 18^, or by detection of specific TLS-associated markers (e.g. CD3, CD20, CD23, DC-LAMP) by immunohistochemistry^6, 18-20^. While informative, these methods are time-consuming, lack scalability, and often miss spatial context.

Recent advancements in digital pathology and artificial intelligence (AI) have enabled the use of deep learning algorithms to detect TLS directly from hematoxylin and eosin (H&E) whole-slide images (WSIs)^21^. Published models leveraging convolutional neural networks (CNNs) have demonstrated high accuracy and scalability in automating TLS identification^22-24^. Despite these developments, the use of different emerging foundation models which are pre-trained architectures using large amount of histology images remains underexplored in detecting TLS. Foundation models offer potential advantages by leveraging excellent transfer learning, enabling feature extraction from diverse datasets prior to a variety of downstream tasks such as tumor detection, biomarker prediction, and survival analysis. Several pathology foundation models use self-supervised learning to enhance image analysis. CTransPath (Swin Transformer)^25^, Phikon (iBOT)^26^, Virchow (DINOv2)^27^, PLIP (Contrastive Learning)^28^, and RETCCL (Clustering-guided Contrastive Learning)^29^ specialize in tasks like biomarker classification, image retrieval, demonstrating significant advancements in histopathology.

In this study (Figure 1), we benchmarked the performance of several commonly used pathology foundation models, including PLIP and Virchow against the pretrained model ResNet50, in identifying TLS within PDAC and HNSCC samples from The Cancer Genome Atlas (TCGA) and RWE datasets.

**Figure 1.**
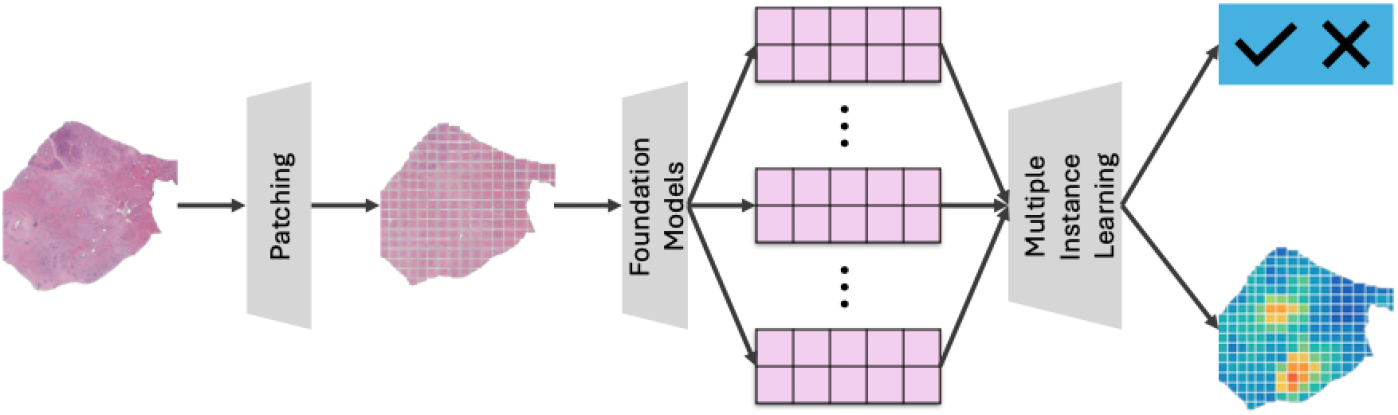
High-level workflow for the histopathology image processing.

By leveraging the strengths of these foundation models in histopathological image analysis, we aim to establish a robust methodology for objective and automated TLS identification and further assess its utility of being a prognostic biomarker associated with clinical outcomes in PDAC and HNSCC.

## Methods

### TCGA PDAC and HNSCC Cohorts

TCGA provides comprehensive genomic profiles for various cancer types, including PDAC and HNSCC. We used 163 patients with high-resolution WSIs and 173 patients with RNA sequencing data, as well as their corresponding clinical information, from a diverse set of pancreatic cancer patients (Table 1.1). Similarly, we selected a subset of the TCGA HNSCC cohort, consisting of detailed clinical information, histopathological images (N=100), and transcriptomic data (N=93) from head and neck cancer patients (Table 1.2). In addition, this study also utilized a HNSCC cohort from a licensed RWE cohort (Tempus AI, Inc., Table 1.2), which comprises longitudinal data from geographically diverse oncology practices. The dataset included 188 samples profiled with histopathology images, of which 185 had matched whole-transcriptome RNA sequencing, as previously described^30^.

**Table 1.1.** Patient characteristics of PDAC-TCGA cohort.

| PDAC-TCGA |  | RNA-seq-based patient group |  | Pathology-based patient group |  |
| --- | --- | --- | --- | --- | --- |
|  |  | TLS High* | TLS Low* | TLS present^ | TLS absent^ |
| Total |  | 84 | 89 | 91 | 72 |
| Gender (%F) |  | 47 (56%) | 31 (35%) | 43 (47%) | 30 (42%) |
| Age |  | 64.9 | 64.5 | 64.5 | 65.3 |
| Pathological Stage | Stage I | 9 | 12 | 12 | 8 |
|  | Stage II | 71 | 72 | 74 | 59 |
|  | Stage III+ | 4 | 3 | 4 | 4 |
|  | Missing | 0 | 0 | 1 | 0 |

**Table 1.2.**
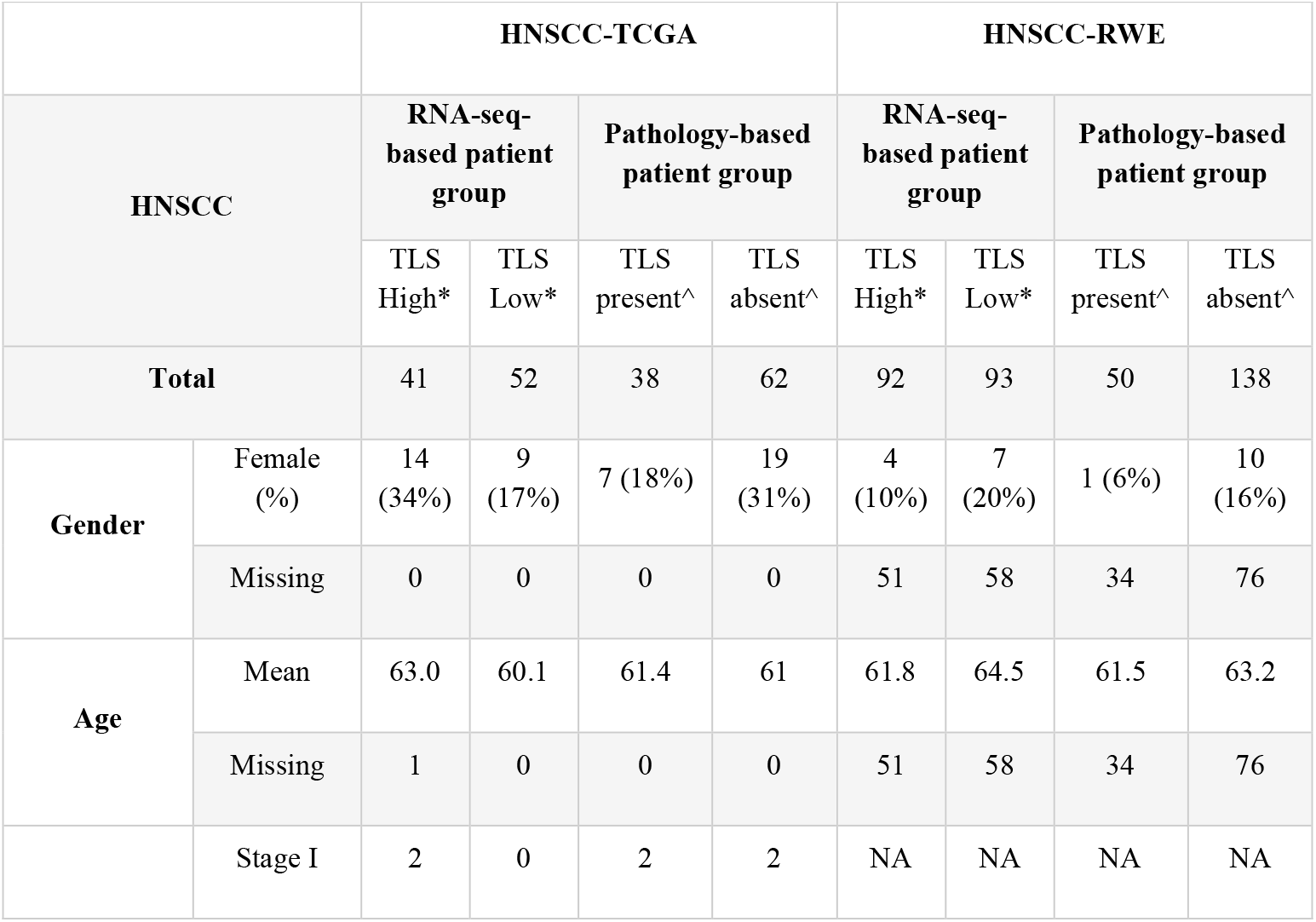
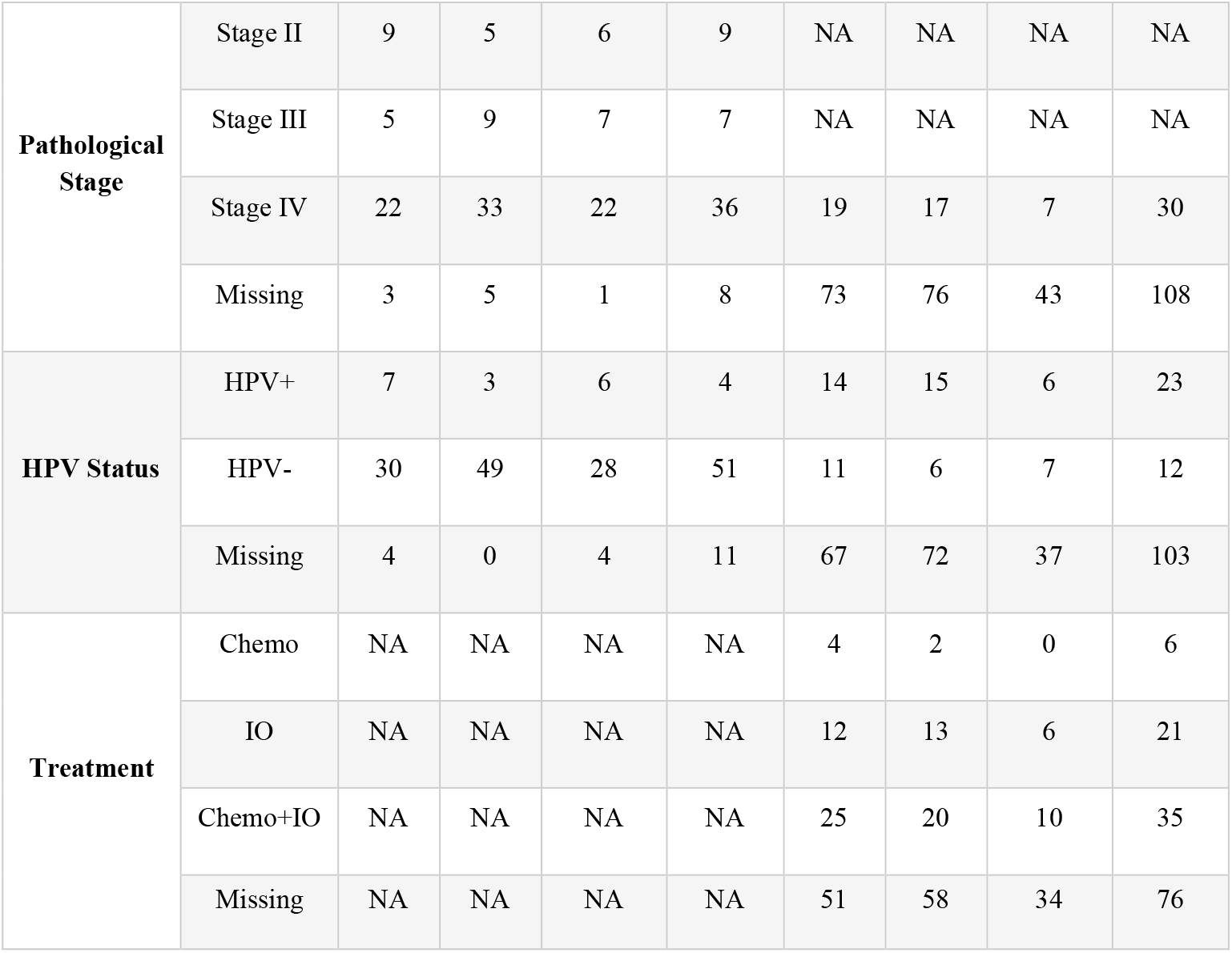
Patient characteristics of HNSCC-TCGA and HNSCC-RWE cohorts.

**Table 2.** Summary of the evaluated pathology foundation models.

| Model | Model Size (Parameters) | Number of Images | Number of Tiles (Millions) | Training Algorithm | Image Source |
| --- | --- | --- | --- | --- | --- |
| **CTransPath** | 28 million | 32,220 | 16 | SRCL | TCGA, PAIP |
| **Phikon** | 86 million | 6,093 | 43 | iBOT | TCGA |
| **Virchow** | 631 million | 1,488,000 | 2,000 | DINOv2 | Memorial Sloan Kettering Cancer Center |
| **PLIP** | Not specified | 208,414 | Not specified | Contrastive Learning | Twitter, LAION |
| **RETCCL** | Not specified | Not specified | Not specified | Contrastive Learning | Custom histopathological datasets |
| **ResNet50** | 25.6 million | 1,280,000 | Not specified | ResNet50 | ImageNet |

### TLS visual assessment on H&E whole slide images

TLS visual assessment on H&E WSIs was performed by Genmab pathologists through Concentriq LS (Proscia, PA, USA). TLS was defined histologically as an organized follicular-like structure of lymphoid cells aggregation of predominant B cell lineage, presence of plasma cells and T cells. The presence of at least one dendritic cell and inclusive or adjacent high endothelial venules were also included as characteristic features of TLS on H&E images^2, 31^. H&E Images with predominant necrosis, significant staining or tissue artefacts and from metastatic lymph node samples were not included for TLS visual assessment. TLS annotation for a subset of images was generated for visualization.

### TLS gene signature calculation and clustering

Gene Set Variation Analysis (GSVA)^32^ scores were calculated for six previously published TLS gene signatures using the normalized bulk transcriptome data (e.g., using TPM or log2-transformed CPM) from TCGA and the RWE cohort. K-means clustering was further applied to the six GSVA scores to derive TLS-high and TLS-low clusters. Uniform manifold approximation and projection (UMAP) algorithm^33^ was used to visualize the clusters.

### Correlation between pathology annotation and TLS signature

Correlation between pathologist-generated TLS label (TLS-present and TLS-absent) and TLS gene signatures-based K-means clusters (TLS-high and TLS-low) were directly evaluated to calculate the concordance score (N _(TLS-present==TLS-high)_ + N _(TLS-absent==TLS-low)_)/N _(Total overlapping samples)_. Welch’s t-test was applied to compare TLS GSVA scores between pathologist-generated TLS-present and TLS-absent groups for both TCGA and RWE cohorts separately for both indications.

### Image processing

Slideflow^34^ package was used for image processing. Tiles were extracted from the whole slide images at 20x magnification and 299 in pixels. During the process, poor quality tiles including background tiles, blurry tiles, or tiles with high whitespace content (>60%), were discarded. Stain normalization was applied using Reinhard algorithm^35^.

### Feature extraction and Multiple-instance learning (ML)

Five different foundation models, including RETCCL, Phikon, CTransPath, PLIP, and Virchow, as well as an ImageNet-trained ResNet50, were used to extract features from image tiles derived from each WSI (Table 2).

The dimensions of the image features ranged from 512 to 2560. The multi-branch clustering-constrained attention multiple instance learning (CLAM_MB) model was used to aggregate and evaluate the effectiveness of the image features. To optimize model performance, a grid search was conducted over a set of hyperparameters, including learning rate, model size, bag size, and batch size. The image set for both indications were split into 75% training and 25% testing. HNSCC TCGA and RWE images were mixed before splitting into training and testing sets. A 3-fold cross-validation method was used to on the training data to evaluate the performance of each hyperparameter combination. The best combination of hyperparameters were run three times to generate 9 data points to have a more accurate estimation. The optimized model was trained on all the training images to predict the outcome in the test data. We utilized several evaluation metrices including accuracy, specificity, recall, area under curve (AUC), and F1 score to measure the model performance.

### Model interpretation with attention heatmaps

Attention heatmaps were generated using the best foundation model in cross-validation to highlight the most relevant regions in predicting the TLS labels. The heatmaps were then further reviewed by a pathologist to evaluate the pathological relevance of the highlighted regions in TLS identification. TLS pathology annotation of the same images was also generated before pathologists reviewed the heatmaps.

## Results

### Patient demographics

A total of 173 pancreatic ductal adenocarcinoma (PDAC) patients from The Cancer Genome Atlas (TCGA) and 288 head and neck squamous cell carcinoma (HNSCC) patients from TCGA (n = 100) and an RWE cohort (n = 188) with pathology assessment were included in this study. Among these, 163 of 173 PDAC patients and 278 of 288 HNSCC patients had matched RNA sequencing (RNAseq) data. No significant differences were observed between TLS signature-based TLS-high and TLS-low groups or pathology-based TLS-present and TLS-absent groups in terms of gender, age, or pathological stage. The majority of PDAC patients were classified as stage 2 (Table 1.1). Similarly, within each HNSCC cohort, no significant differences were detected in patient characteristics, including gender, age, disease stage, human papillomavirus (HPV) status, or treatment history, between TLS groups (signature-based or pathology-based).

Additionally, demographic characteristics, including gender, age, and disease stage, were comparable between the HNSCC-TCGA and HNSCC-RWE cohorts (Table 1.2). However, most HNSCC-TCGA patients were immunotherapy (IO) naïve, whereas the majority of HNSCC-RWE patients had received prior treatment. Furthermore, HPV positivity was lower in the HNSCC-TCGA cohort (11.24%) compared to the HNSCC-RWE cohort (60.4%).

### TLS label generation and evaluation

We first generated GSVA scores using transcriptomic data from TCGA and the RWE cohort for the six published TLS signatures (Supplementary Table 1). A K-means clustering algorithm was applied to classify TLS-high and TLS-low groups (Figure 2) for PDAC, HNSCC-TCGA, and HNSCC-RWE separately. For PDAC, we identified 84 TLS-high and 89 TLS-low slides, and for HNSCC, we identified 133 TLS-high (41 TCGA + 92 RWE) and 145 TLS-low (52 TCGA + 93 RWE) slides. Separately, pathologists performed a pathological review on H&E images to classify 91 TLS-present and 72 TLS-absent slides for PDAC, and 88 TLS-present (38 TCGA + 50 RWE) and 200 TLS-absent (62 TCGA + 138 RWE) for HNSCC. Notably, significant variations of TLS histological features between PDAC and HNSCC H&E WSIs are appreciated, such as the size, number of immune cells, cell density, maturation status, and spatial distribution in TME on H&E images. Concordance between pathology-assessed TLS levels and K-means clustering was 56.6% for PDAC and 41.0% for HNSCC (54.9% TCGA, 34.06% RWE, Supplementary Figure 1).

**Figure 2.**
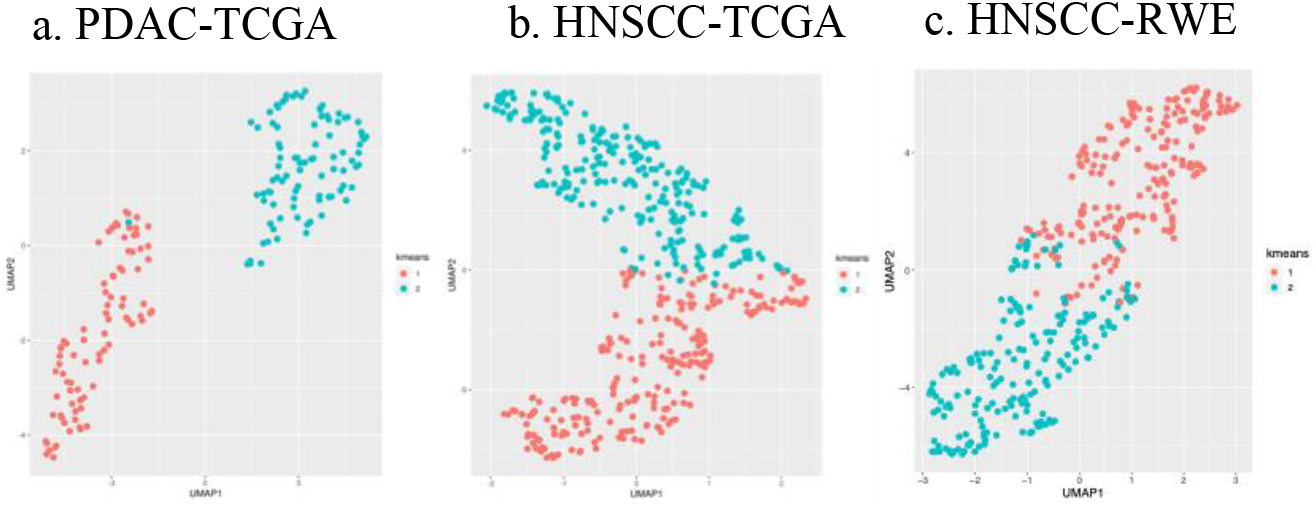
Patient clustering with TLS signatures in PDAC-TCGA (a), HNSCC-TCGA (b), and HNSCC-RWE (c).

The lower concordance rate in HNSCC was observed primarily in the RWE cohort, which exhibited greater heterogeneity in the images. To further explore the correlation between pathological review and individual TLS signatures, we conducted Welch’s t-test to compare TLS signature levels between TLS-present and TLS-absent groups in both PDAC and HNSCC (Supplementary Figure 2). In PDAC, TLS-present cases showed consistently higher TLS signature levels (p<0.05) in 5 out of 6 different TLS signatures (Supplementary Figure 2). In the HNSCC cohort, TLS-present tumors similarly displayed significantly higher TLS signature levels (p<0.05) in 5 out of 6 signatures in the HNSCC-RWE cohort. However, none of the comparisons in HNSCC-TCGA reached significance, despite showing similar trends (Supplementary Figure 2).

### TLS Prediction in PDAC

The performance of six feature extractors—ResNet50, Phikon, RETCCL, CTransPath, PLIP, and Virchow—on predicting both RNAseq-based and pathology-derived TLS labels in the PDAC cohort is presented in Figure 3. Overall, all models struggled to predict RNAseq-based labels, with average AUCs ranging from 0.54 to 0.65 using 3x3 cross-validation (Figure 3.a; Supplementary table 2). In contrast, all models performed better in predicting pathology labels with AUCs ranging from 0.71 to 0.84 with 3x3 cross-validation (Figure 3.a; Supplementary table 3). PLIP achieved highest AUC of 0.84 (±0.06), followed by CTransPath with an AUC of 0.78 (±0.06) on the training data (Figure 3.a; Supplementary table 3). This trend persisted in the testing data, where all models demonstrated improved performance on pathology labels (AUC: 0.64–0.94) compared to K-means (AUC: 0.40–0.63) (Figure 3.b; Table 3). PLIP exhibited particularly robust performance on the testing data, with an AUC of 0.94 and an F1 score of 0.95 when predicting pathology labels. CTransPath closely followed, achieving an AUC of 0.89 and an F1 score of 0.91. Surprisingly, ImageNet-trained ResNet50 also showed respectable performance, with an AUC of 0.85 and an F1 score of 0.86. The best-performing feature extractor for K-means predictions on testing data was ImageNet-pretrained ResNet50, with an AUC of 0.63 and an F1 score of 0.67 (Figure 3.b; Table 3).

**Table 3.**
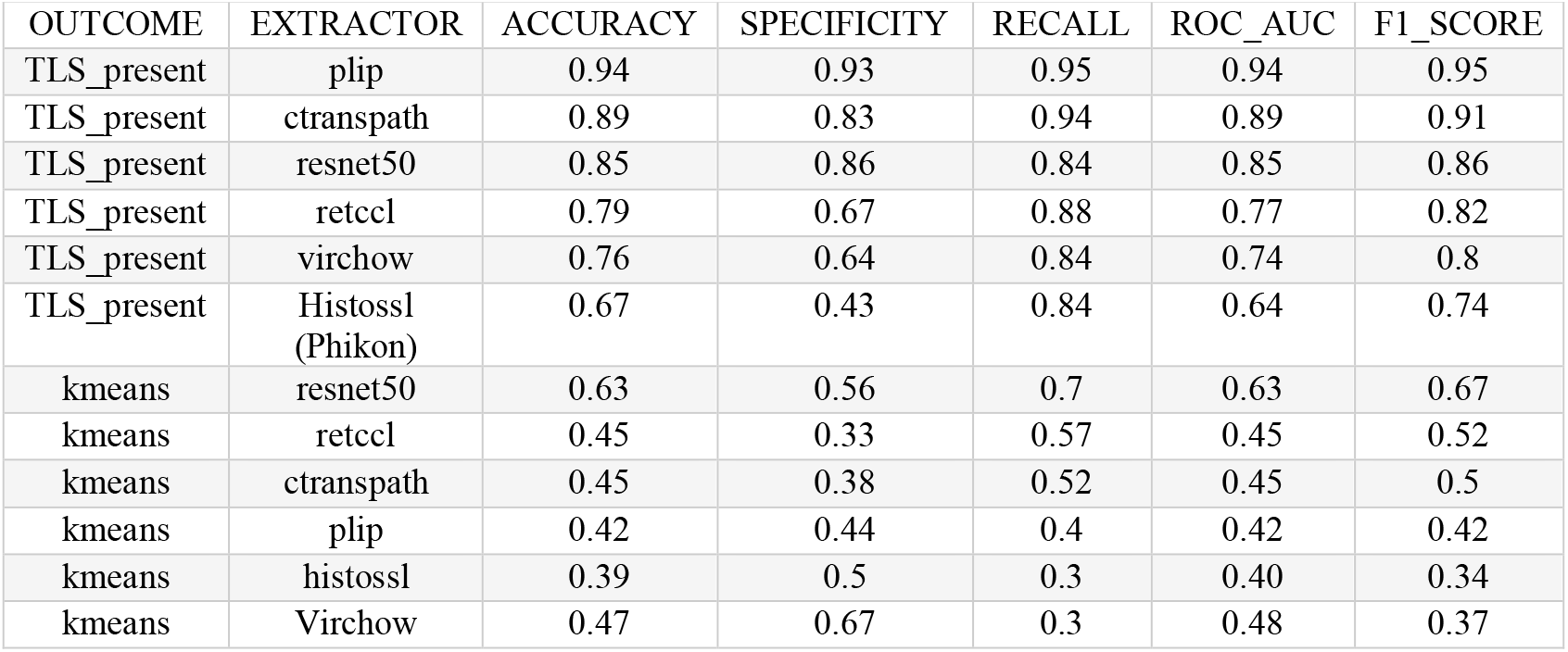
Detailed performance of foundation models in predicting TLS in PDAC test data.

**Figure 3.**
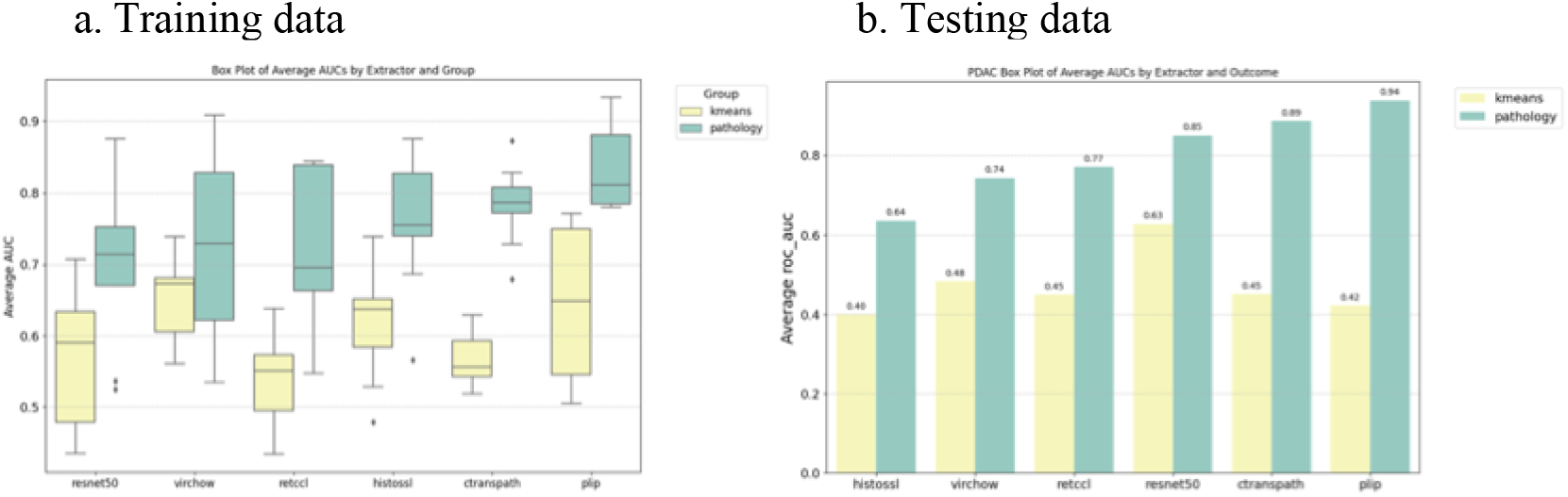
Performance of foundation models in predicting TLS in PDAC of (a) cross validation (3x3) of training set and (b) testing set.

### TLS Prediction in HNSCC

The same feature extractors were evaluated in the HNSCC cohort for predicting both RNAseq-based and pathology-derived TLS status (Figure 4). Like PDAC, all models struggled to predict RNAseq-based labels, with average AUCs ranging from 0.63 to 0.70 in 3x3 cross-validation (Figure 4.a; Supplementary table 4). Most models again demonstrated better performance on pathology labels, except for ResNet50 (0.59 (±0.05)) (Figure 4.a; Supplementary table 5). CTransPath achieved the highest performance, with an AUC of 0.80 (±0.05), followed by Virchow and RETCCL with an AUC of 0.74 (±0.02) and 0.74 (±0.09) respectively (Figure 4.a; Supplementary table 5). In testing data, however, all models struggled with both pathology and K-means predictions, with Virchow achieving the highest AUC of 0.71 and an F1 score of 0.59 for pathology label predictions (Figure 4.b; Table 5). Notably, ResNet50 performed poorly on the HNSCC pathology label with an AUC of 0.48 and an F1 score of 0.35, a marked contrast to its robust performance in PDAC (Figure 4.b; Table 5). The significant drop in pathology label prediction accuracy in HNSCC compared to PDAC may be attributable to a more complex TME, greater heterogeneity in tumor location based on internal pathology assessment and previous publications^36, 37^, and the mixed dataset from TCGA and the RWE cohort.

**Table 4.** Detailed performance of foundation models in predicting TLS in HNSCC test data.

| OUTCOME | EXTRACTOR | ACCURACY | SPECIFICITY | RECALL | ROC_AUC | F1_SCORE |
| --- | --- | --- | --- | --- | --- | --- |
| TLS_present | virchow | 0.72 | 0.76 | 0.68 | 0.71 | 0.59 |
| TLS_present | plip | 0.7 | 0.71 | 0.68 | 0.70 | 0.58 |
| TLS_present | ctranspath | 0.72 | 0.76 | 0.63 | 0.69 | 0.57 |
| TLS_present | retccl | 0.64 | 0.64 | 0.63 | 0.64 | 0.51 |
| TLS_present | Histossl (Phikon) | 0.61 | 0.6 | 0.63 | 0.62 | 0.49 |
| TLS_present | resnet50 | 0.48 | 0.49 | 0.47 | 0.48 | 0.35 |
| kmeans | virchow | 0.56 | 0.41 | 0.56 | 0.56 | 0.58 |
| kmeans | retccl | 0.6 | 0.48 | 0.71 | 0.59 | 0.66 |
| kmeans | plip | 0.59 | 0.45 | 0.71 | 0.58 | 0.65 |
| kmeans | Histossl (Phikon) | 0.59 | 0.55 | 0.62 | 0.58 | 0.62 |
| kmeans | ctranspath | 0.48 | 0.41 | 0.53 | 0.47 | 0.52 |
| kmeans | resnet50 | 0.51 | 0.55 | 0.47 | 0.51 | 0.51 |

**Figure 4.**
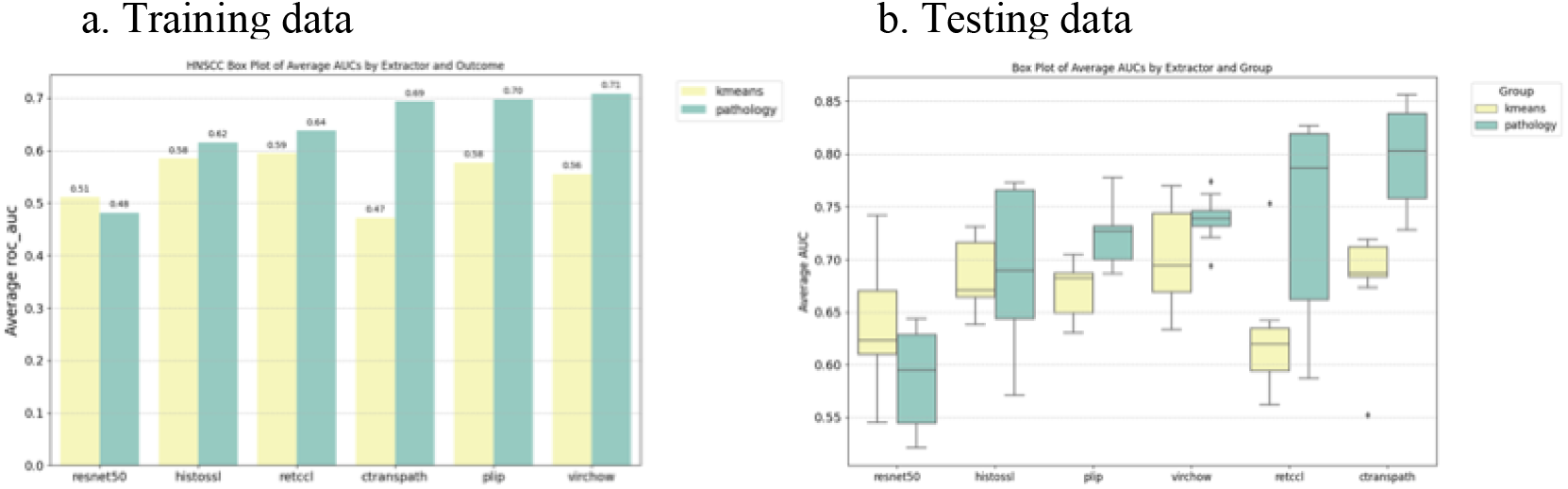
Performance of foundation models in predicting TLS in HNSCC of (a) cross validation (3x3) of training set and (b) testing set.

### Model interpretation

To better understand how the model detected the TLS and explore the disparities in model performance, pathology annotation and attention heatmaps were generated for both PDAC and HNSCC cohorts (Figure 5). Attention heatmaps highlight regions of histopathology slides that contribute most to the model’s prediction, offering crucial interpretability in digital pathology by allowing pathologists to assess whether the AI is focusing on biologically and clinically relevant tissue features. Attention heatmaps generated by the CLAM on PLIP features for pathology-based TLS status in PDAC showed high-level of consistency with pathology annotation (Figure 5.a). In contrast, attention heatmaps generated with Virchow extracted features for HNSCC struggled to focus on TLS regions-only (Figure 5.b). Pathological evaluation indicated that the high-attention tiles identified by the model in HNSCC were influenced by pre-existing lymph nodes and non-TLS immune cell infiltration, which may exhibit visual similarities to TLS.

**Figure 5.**
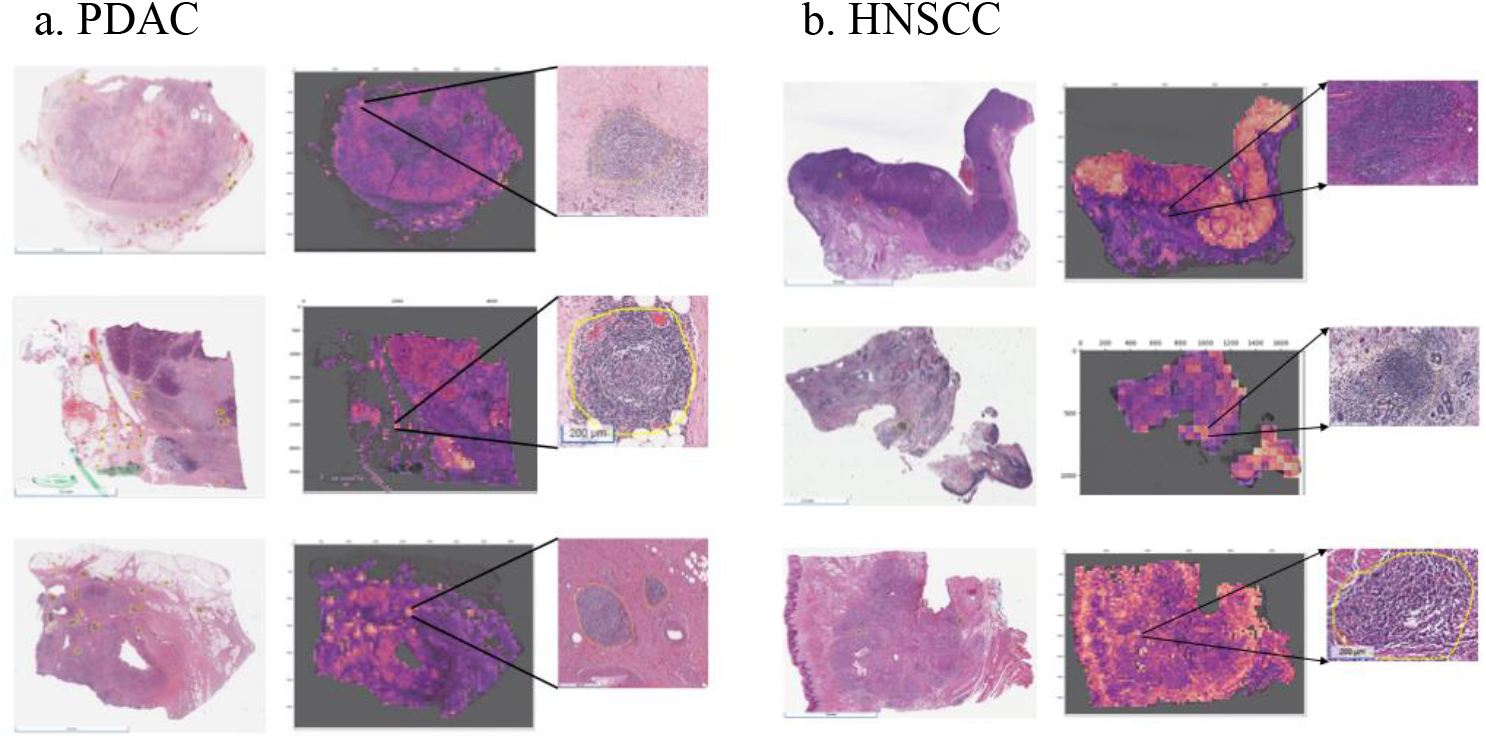
Attention maps of PDAC (a) and HNSCC (b) slides with TLS.

## Discussion

Self-supervised learning (SSL) has significantly advanced computational pathology by enabling the training of foundation models on large pathology image datasets. The increasing availability of publicly released models by both academic and private institutions is fostering innovation in the next generation of predictive pathology tools. Foundation models, particularly in weakly-supervised algorithms, have demonstrated superior performance and generalizability compared to traditional supervised methods. These models have been instrumental in cancer research, facilitating tumor diagnosis, biomarker prediction, and prognostic assessments^38, 39^. Since 2022, foundation models have been integrated into weakly supervised pipelines, leading to improved diagnostic accuracy and broader applicability. As more foundation models are trained, benchmarking studies on clinically relevant tasks have also become available. Campanella et al. benchmarked eight foundation models on nine disease detection and eleven biomarker prediction tasks, finding that while newer models generally outperformed ImageNet-pretrained encoders and CTransPath, their performance varied by task and was not significantly impacted by model size^38^. Similarly, Neidlinger et al. demonstrated that model performance varied by task, with the quality and cleanliness of training data outweighing data volume or the training algorithm^39^.

In this study, we evaluated the utility of the pathology foundation models for a unique task— detecting TLS presence, defined by morphological features or transcriptomic gene signatures, in two distinct indications: PDAC and HNSCC. As previously shown, detecting TLS via computational methods typically requires extensive annotation and segmentation efforts^21-24^.

Here, we demonstrate that pathology foundation models can serve as an effective and out-of-the-box alternative for detecting morphologically defined TLS in tumor types with less complex TME. Consistent with Campanella et al. and Neidlinger et al.^38, 39^, we also found that model performance varies by difficulty of the task while model size was not a significant factor.

In general, pathology foundation models outperformed the ImageNet-pretrained ResNet50 in both the PDAC and HNSCC cohorts when predicting morphology-based TLS status. Although the foundation models consistently outperformed ResNet50 in HNSCC slide images, they all achieved lower mean AUC scores compared to their performance on PDAC tumors. This disparity is likely due to the heterogeneous tumor microenvironment of HNSCC, which may cause the foundation models to struggle in accurately identifying specific factors among multiple complex features. For example, pre-existing lymph nodes were frequently present in the HNSCC TME and may have affected the model’s performance.

The largest foundation model evaluated, Virchow, did not demonstrate superior performance over smaller models, despite narrowly achieving the highest performance on the HNSCC testing data. This aligns with previous findings that model size does not necessarily correlate with performance.

When predicting pathology-based TLS in PDAC, ResNet50 achieved a comparable AUC to most pathology foundation models in the training set and ranked third in the testing set. Interestingly, in the HNSCC cohort, the gap between pathology foundation models and ResNet50 widened, with foundation models consistently outperforming ResNet50 in predicting pathology-based TLS. These findings suggest that while pathology image-pretrained models still have room for improvement, they provide the most value in pathologically challenging tasks.

Contrary to pathology-based label prediction, all pathology models performed poorly and did not outperform ResNet50 in identifying signature-based TLS status. This could suggest that while the foundation models excel in H&E image recognition capabilities, they struggle in RNA-based biomarker predictions, highlighting a potential limitation in their versatility across modalities.

This may be partly due to the fact that TLS gene signatures are not necessarily TLS-specific and may not directly correlate with morphological features visible on H&E-stained slides. Given that current pathology foundation models largely rely on morphological features, incorporating omics data into their training could improve predictive performance, especially when target labels are defined by molecular or transcriptomic signatures^40^.

Overall, model performance varied by task complexity rather than size, aligning with recent benchmarking findings. One limitation of this study is that only five foundation models were evaluated, each pretrained using different image sizes and algorithms. This selection was not intended to provide an exhaustive benchmark of all available models. Rather, the aim was to illustrate the practical utility and inherent limitations of representative models within the context of this specific application.

## Conclusion

This study demonstrated that pathology foundation models effectively detected TLS presence in PDAC and HNSCC, outperforming ImageNet-pretrained ResNet50 in morphology-based predictions. However, foundation models struggled with transcriptomic signature-based TLS prediction, revealing cross-modal generalization challenges, and performed worse in heterogeneous HNSCC microenvironments, highlighting limitations in complex pathological contexts. Despite these challenges, pathology foundation models show strong potential as the backbone of automated digital pathology pipelines for task-specific applications. The widespread availability and standardization of H&E-stained images make them a compelling source for model deployment and scaling, particularly in biomarker discovery and patient stratification. Continued refinement will be essential to enhance their versatility and robustness across modalities and tumor heterogeneity.

## Data availability

TCGA: https://portal.gdc.cancer.gov/.

### Commercial cohort

Deidentified data used in the research were collected in a real-world health care setting and are subject to controlled access for privacy and proprietary reasons. When possible, derived data supporting the findings of this study have been made available within the paper and its Supplementary Figures/Tables. Restrictions apply to the availability of additional data, which were used under license for this study.

## Consents and Ethics

### Consent

This study was conducted on de-identified health information subject to an IRB exempt determination (Advarra Pro00072742) and did not involve human subject research.

### Funding Information

This study was sponsored by Genmab.

