## Supplementary Material for "Comparative Analysis of Pathology Foundation Models for Automated Detection of Tertiary Lymphoid Structures in H&E-Stained Digital Pathology Images"

^3^Cornell University, Ithaca, New York

**Supplementary Material**

Supplementary table 1. TLS gene signatures

| TLS_1 | CD79B | EIF1AY | PTGDS | RBP5 | CCR6 | SKAP1 | LAT | CETP | CD1D |  |  |  |
| --- | --- | --- | --- | --- | --- | --- | --- | --- | --- | --- | --- | --- |
| TLS_2 | CCL19 | CCL21 | CXCL13 | CCR7 | CXCR5 | SELL | LAMP3 |  |  |  |  |  |
| TLS_3 | FCRL5 | IDO1 | IFNG | BTLA |  |  |  |  |  |  |  |  |
| Chemokine signature | CCL2 | CCL3 | CCL4 | CCL5 | CCL8 | CCL18 | CCL19 | CCL21 | CXCL9 | CXCL10 | CXCL11 | CXCL13 |
| TFH signature | CXCL13 | CD200 | FBLN7 | ICOS | SGPP2 | SH2D1A | TIGIT | PDCD1 |  |  |  |  |
| TH1 cell and B cell signature | CD4 | CCR5 | CXCR3 | CSF2 | IGSF6 | IL2RA | CD38 | CD40 | CD5 | MS4A1 | SDC1 | GFI1 |
|  | IL11 | IL1R2 | IL10 | CCL20 | IRF4 | TRAF6 | STAT5A |  |  |  |  |  |

Supplementary table 2. Model hyperparameters for kmeans prediction in PDAC training data

| EXTRACTOR | LR | BAG_SIZE | EPOCHS | BATCH_SIZE | MODEL | MODEL_SIZE | BAG_WEIGHT | DROPOUT | OPT | AVERAGE_AUCS | SD_AUCS |
| --- | --- | --- | --- | --- | --- | --- | --- | --- | --- | --- | --- |
| virchow | 0.0001 | 1024 | 24 | 1 | clam_mb | big | 0.7 | 0.1 | adam | 0.65 | 0.05 |
| plip | 0.001 | 512 | 24 | 1 | clam_mb | big | 0.7 | 0.1 | adam | 0.65 | 0.11 |
| Histossl (Phikon) | 0.0001 | 512 | 24 | 1 | clam_mb | big | 0.7 | 0.1 | adam | 0.62 | 0.08 |
| resnet50 | 0.0001 | 512 | 24 | 1 | clam_mb | big | 0.7 | 0.1 | adam | 0.58 | 0.10 |
| ctranspath | 0.0001 | 512 | 24 | 1 | clam_mb | big | 0.7 | 0.1 | adam | 0.57 | 0.04 |
| retccl | 0.01 | 512 | 24 | 1 | clam_mb | small | 0.7 | 0.1 | adam | 0.54 | 0.07 |

Supplementary table 3. Model hyperparameters for pathology TLS prediction in PDAC training data

| EXTRACTOR | LR | BAG_SIZE | EPOCHS | BATCH_SIZE | MODEL | MODEL_SIZE | BAG_WEIGHT | DROPOUT | OPT | AVERAGE_AUCS | SD_AUCS |
| --- | --- | --- | --- | --- | --- | --- | --- | --- | --- | --- | --- |
| plip | 0.0001 | 512 | 24 | 1 | clam_mb | big | 0.7 | 0.1 | adam | 0.84 | 0.06 |
| ctranspath | 0.0001 | 512 | 24 | 1 | clam_mb | big | 0.7 | 0.1 | adam | 0.78 | 0.06 |
| Histossl (Phikon) | 0.001 | 512 | 24 | 1 | clam_mb | big | 0.7 | 0.1 | adam | 0.76 | 0.1 |
| retccl | 0.01 | 2048 | 24 | 1 | clam_mb | big | 0.7 | 0.1 | adam | 0.72 | 0.1 |
| virchow | 0.01 | 512 | 24 | 1 | clam_mb | small | 0.7 | 0.1 | adam | 0.72 | 0.13 |
| resnet50 | 0.01 | 2048 | 24 | 1 | clam_mb | multiscale | 0.7 | 0.1 | adam | 0.71 | 0.12 |

Supplementary table 4. Model hyperparameters for kmeans prediction in HNSCC training data

| EXTRACTOR | LR | BAG_SIZE | EPOCHS | BATCH_SIZE | MODEL | MODEL_SIZE | BAG_WEIGHT | DROPOUT | OPT | AVERAGE_AUCS  ($\boldsymbol{\pm}\mathbf{SD}$) |
| --- | --- | --- | --- | --- | --- | --- | --- | --- | --- | --- |
| ctranspath | 0.01 | 2048 | 24 | 1 | Clam_mb | small | 0.7 | 0.1 | adam | 0.68 (±0.05) |
| retccl | 0.01 | 1024 | 24 | 1 | clam_mb | multiscale | 0.7 | 0.1 | adam | 0.63 (±0.05) |
| plip | 0.0001 | 2048 | 24 | 1 | clam_mb | multiscale | 0.7 | 0.1 | adam | 0.67 (±0.03) |
| Histossl (Phikon) | 0.0001 | 512 | 24 | 1 | clam_mb | small | 0.7 | 0.1 | adam | 0.68 (±0.04) |
| resnet50 | 0.001 | 512 | 24 | 1 | clam_mb | big | 0.7 | 0.1 | adam | 0.64 (±0.06) |
| virchow | 0.0001 | 1024 | 24 | 1 | clam_mb | multiscale | 0.7 | 0.1 | adam | 0.70 (±0.05) |

Supplementary table 5. Model hyperparameters for pathology TLS prediction in HNSCC training data

| EXTRACTOR | LR | BAG_SIZE | EPOCHS | BATCH_SIZE | MODEL | MODEL_SIZE | BAG_WEIGHT | DROPOUT | OPT | AVERAGE_AUCS  ($\boldsymbol{\pm}\mathbf{SD}$) |
| --- | --- | --- | --- | --- | --- | --- | --- | --- | --- | --- |
| ctranspath | 0.001 | 512 | 24 | 1 | Clam_mb | Multiscale | 0.7 | 0.1 | adam | 0.80 ($\pm$0.05) |
| retccl | 0.01 | 512 | 24 | 1 | clam_mb | multiscale | 0.7 | 0.1 | adam | 0.74 ($\pm$0.09) |
| plip | 0.0001 | 512 | 24 | 1 | clam_mb | small | 0.7 | 0.1 | adam | 0.72 ($\pm$0.03) |
| Histossl (Phikon) | 0.001 | 512 | 24 | 1 | clam_mb | multiscale | 0.7 | 0.1 | adam | 0.69 ($\pm$0.08) |
| resnet50 | 0.01 | 512 | 24 | 1 | clam_mb | big | 0.7 | 0.1 | adam | 0.59 ($\pm$0.05) |
| virchow | 0.0001 | 512 | 24 | 1 | clam_mb | small | 0.7 | 0.1 | adam | 0.74 ($\pm$0.02) |

Supplementary Table 6. Tile numbers

| Tile number (N) | Pathology Label | K-means |
| --- | --- | --- |
| PDAC | 354299 | 368312 |
| HNSCC | 289832 | 274198 |

Supplementary Figure 1. Concordance between pathology label and gene signature-based K-means

a. PDAC

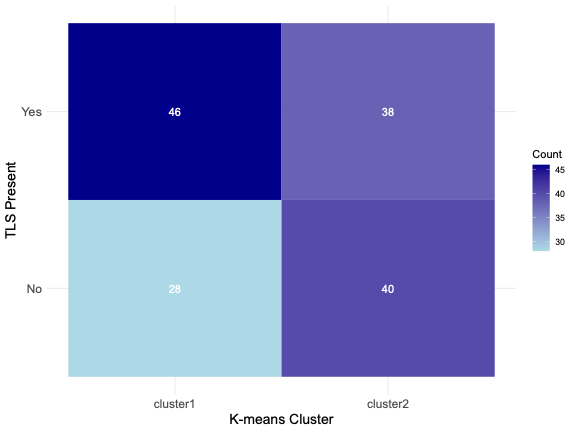

b. HNSCC

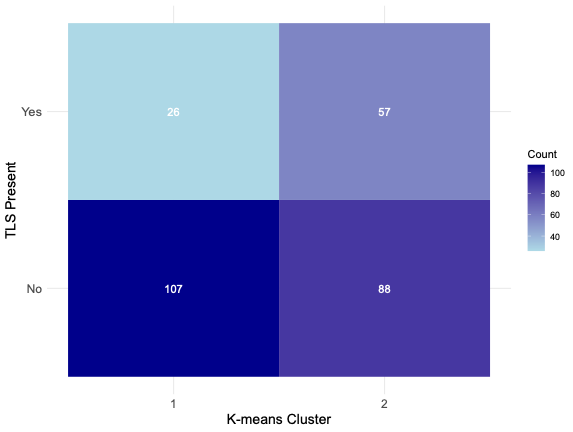

Supplementary Figure 2. Correlation of pathology annotation and TLS signatures

a. HNSCC-RWE

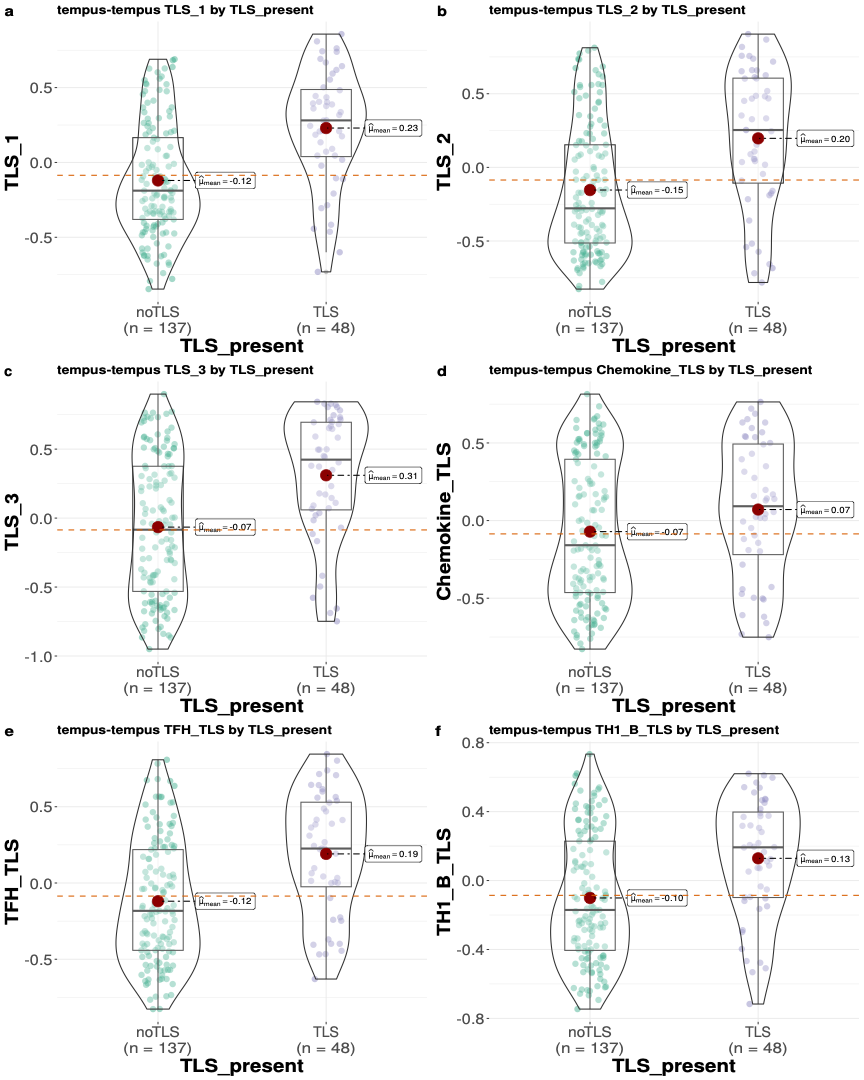

b. HNSCC-TCGA

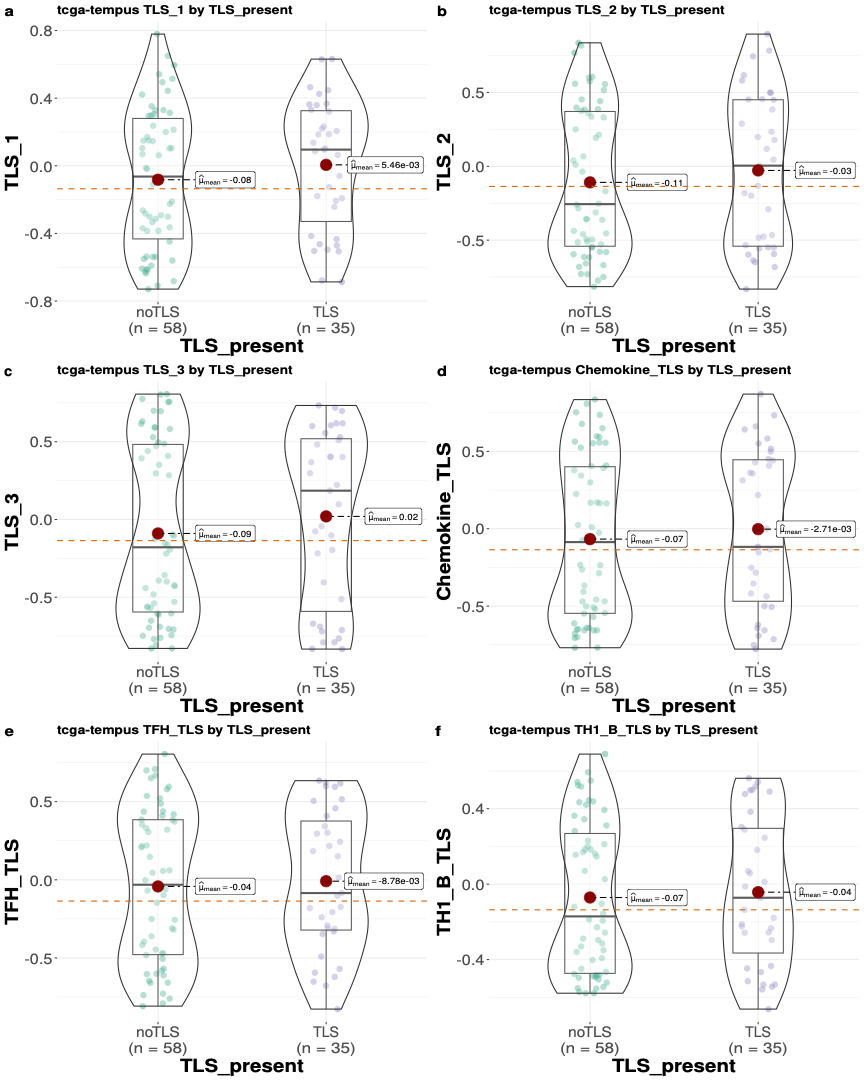

c. PDAC-TCGA

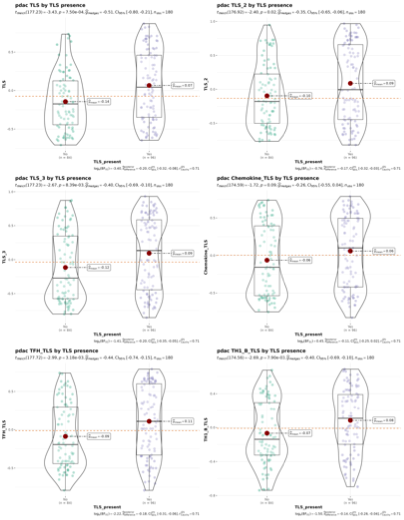
